# Difference contact maps: from what to why in the analysis of the conformational flexibility of proteins

**DOI:** 10.1101/575696

**Authors:** Mallika Iyer, Zhanwen Li, Lukasz Jaroszewski, Mayy Sedova, Adam Godzik

**Affiliations:** Graduate School of Biomedical Sciences, Sanford Burnham Prebys Medical Discovery Institute, 10901 North Torrey Pines Road, La Jolla, CA, 92037, USA; Biosciences Division, University of California Riverside School of Medicine, 900 University Ave., Riverside, CA, 92521, USA

## Abstract

Protein structures, usually visualized in various highly idealized forms focusing on the three-dimensional arrangements of secondary structure elements, can also be described as lists of interacting residues or atoms and visualized as two-dimensional distance or contact maps. We show that contact maps provide an ideal tool to describe and analyze differences between structures of proteins in different conformations. Expanding functionality of the PDBFlex server and database developed previously in our group, we describe how analysis of difference contact maps (DCMs) can be used to identify critical interactions stabilizing alternative protein conformations, recognize residues and positions controlling protein functions and build hypotheses as to molecular mechanisms of disease mutations.

**Author Summary:** Protein folding is driven, and the three-dimensional structures of proteins are defined, by the non-local interactions between amino acid chains. However, the usual visualizations of protein structures focus on the overall chain topology and do not provide much information about the interactions defining them. Here we explore contact map visualization of protein structures, which directly focuses on interactions stabilizing the structure and thus can be easily used to understand effects of mutations on protein structure. We show that contact map visualization is particularly useful for comparing alternative conformations of proteins and identifying critical positions controlling functional flexibility.

## Introduction

Protein structures have complex three-dimensional shapes and are most often visualized as cartoons depicting their overall arrangement of secondary structure elements and neglecting interaction details. Such cartoons were popularized by Jane Richardson [2] and gained wide popularity thanks to programs such as PyMol [3] (see Figure 1A). Other visualization styles: topology diagrams [4], distance [5] or contact [6] maps are occasionally used, but their popularity doesn’t compare to that of ribbon diagrams, which became de facto standards in presenting protein structures in manuscripts and books.

**Figure 1.**
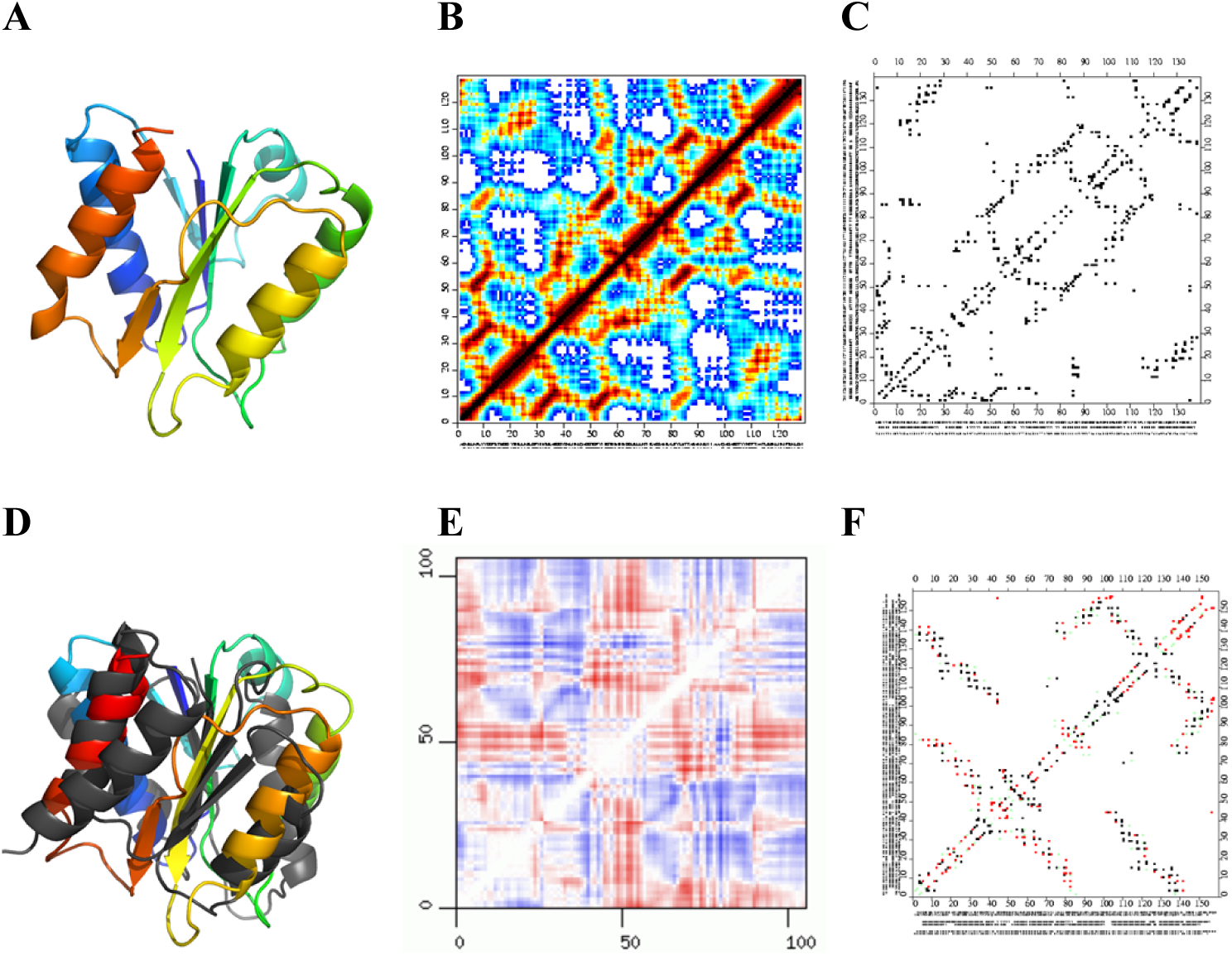
Examples of visualization of protein structures A) ribbon diagram B) distance and C) contact map and differences between them D) overlap of two structures E) difference distance map and F) difference contact map

Widespread use of such images to depict protein structures, often combined with wording that talks about “the structure” when referring to entities illustrated by such images, reinforced the misconception that protein structures are unique and static. However, protein structures are far from static and, as any physical system in constant temperature, can assume any of the conformations from the canonical ensemble describing the system [7]. Protein functions often include cycling through various functional isoforms that correspond to different neighborhoods in the conformational ensemble. For many proteins, single conformations representing different functional forms have been captured experimentally and are available as different coordinate sets for the same protein in the Protein Data Bank (PDB) [8]. Differences between such alternative conformations are difficult to illustrate by ribbon diagrams and are often described verbally or shown in detail only for the most relevant, but small section of the structure, such as for instance an active site.

The most often used measure of structural difference between protein structures is the root mean square deviations (RMSD) between Cα atoms [9]. While useful for classifying and rank ordering of (dis-)similarity of protein structures, it is a global measure that doesn’t give much information about the details of the differences and treats on equal footing a protein pair with significant, but localized differences in one loop with a pair with subtle, but distributed differences. Similar to other popular protein structure similarity/dissimilarity measures, such as TM-score [10], RMSD is useful for identifying the most similar (or divergent) structures from a group, but not to describe the details nor mechanisms of the divergence.

Protein structure visualizations that directly focus on interactions stabilizing it may be better suited for this purpose and were indeed quite popular in the early days of structural biology, but mostly fell out of favor with the growing popularity of ribbon diagrams. For instance, difference distance (Figure 1B) or contact (Figure 1C) maps can be used to compare protein structures and analyze the details of differences between functional states [11] (See Figure 1E and 1F, respectively). But as historically the main focus of structural biology was the exploration of the protein universe, classification and initial characterization of novel structures was a priority and new tools and visualizations became popular. Now structural biology is increasingly focusing on details of protein function rather than on initial structural characterization of novel proteins and we are in need of tools for the in depth analysis of differences between different conformations of otherwise similar or identical proteins. An increasing percentage of protein coordinate sets deposited to the PDB consists of multiple depositions of independently solved structures of the same protein, either in different functional states or simply in different conditions, resolution etc., resulting in apparently redundant deposition. For instance, in the latest PDB release (data from Jan 25, 2019) there were 478,448 independent protein chains with 70,567 unique (at 100% identity level) sequences – for an over 6-fold redundancy (this would further fall to 56,973 i.e. 8.4-fold redundancy at the 95% threshold similarity level, used in this project).

Many global analyses of structures in the Protein Data Bank try to remove this redundancy, selecting a single chain to represent each group of alternative conformations and both specialized servers [12] and the PDB itself provide lists such of such non-redundant structures. This approach, perhaps inadvertently, reinforces the unique structure paradigm by treating all information about structural variation within a conformational ensemble of individual proteins as noise to be removed from analysis. This surprisingly large variation was studied previously in our group [13] and to study it further, we developed PDBFlex [14], a server that illustrates and performs automated analysis of all clusters of coordinates sets for almost-identical (95% sequence identity) proteins. However, PDBFlex, as well as other similar resources [15], is based almost entirely on using the language of RMSD and global comparisons, thus focusing on the question of classifying and describing the structural divergence within the cluster – the “what” question. Here we would like to approach the “why” question – which interactions define the structural variations, how it can be influenced by mutations, etc. We believe that to answer these questions we need to get away from global descriptions such as RMSD and develop a new language to describe the differences between protein conformations, a language anchored in interactions.

## Results

We decided to characterize the differences between two conformations of a protein in a language of residue interactions and analyze differences between contact maps of proteins. Such an approach was used in the past by us and other groups to optimize protein structural alignment [6], to classify protein structures [16] or as a general tool for protein structure comparison [17]. Finding an optimal overlap between protein contact maps became a popular algorithmic problem [18]. We hypothesize that such analysis would allow us to identify residues that differentially stabilize alternative conformations (**D**ifferentially **S**tabilizing **R**esidue**s** or DSRs). We hypothesize that DSR-based description of differences between alternative protein conformations, in contrast to more popular RMSD-based descriptions, is a natural way to identify factors that may control the natural balance between functional/structural states of a protein and which, if modified by disease mutations or interactions with drugs, may disrupt this balance.

To identify the possible DSRs and evaluate this approach on a large set of proteins, we compared residue contact maps for conformations representing the most divergent structure models of all proteins with “redundant” structures deposited in the Protein Data Bank [8]. The starting point for this approach was the PDBFlex [14] database developed previously in our lab.

### Non-redundant structures in the PDB illustrate the scope of conformational differences between alternative protein conformations

The PDBFlex database is based on clusters of PDB entries with >95% sequence identity, representing independently solved structural models of the same protein, with the 95% sequence identity threshold adapted to allow for variations in construct boundaries and/or single mutations. From each cluster, the two sets of coordinates with largest RMSD were selected to represent the two most divergent conformations of the protein in the cluster, although this approach can be applied to study the differences between any two conformations. Figure 2 shows a histogram of maximum RMSD for all clusters in the PDBFlex database (snapshot from 30 April 2018). The main peak around 0.4Å RMSD (9079 clusters with RMSD ≤ 0.5 Å) shows that for most clusters the structural differences are small. We hypothesize that such fluctuations describe movements around the mean structure, like in a short molecular dynamics simulation [19] or protein normal mode analysis [20]. However, the distribution also shows a long tail towards higher RMSDs (20934 clusters with RMSD > 0.5 Å), which correspond to significant structural differences, such as the difference between the active and inactive conformation of BRAF (6.2 Å RMSD) and the open and closed forms of adenylate kinase (7.3 Å RMSD). This tail is seen more clearly when the y-axis is log-scaled (Figure 2 inset).

**Figure 2.**
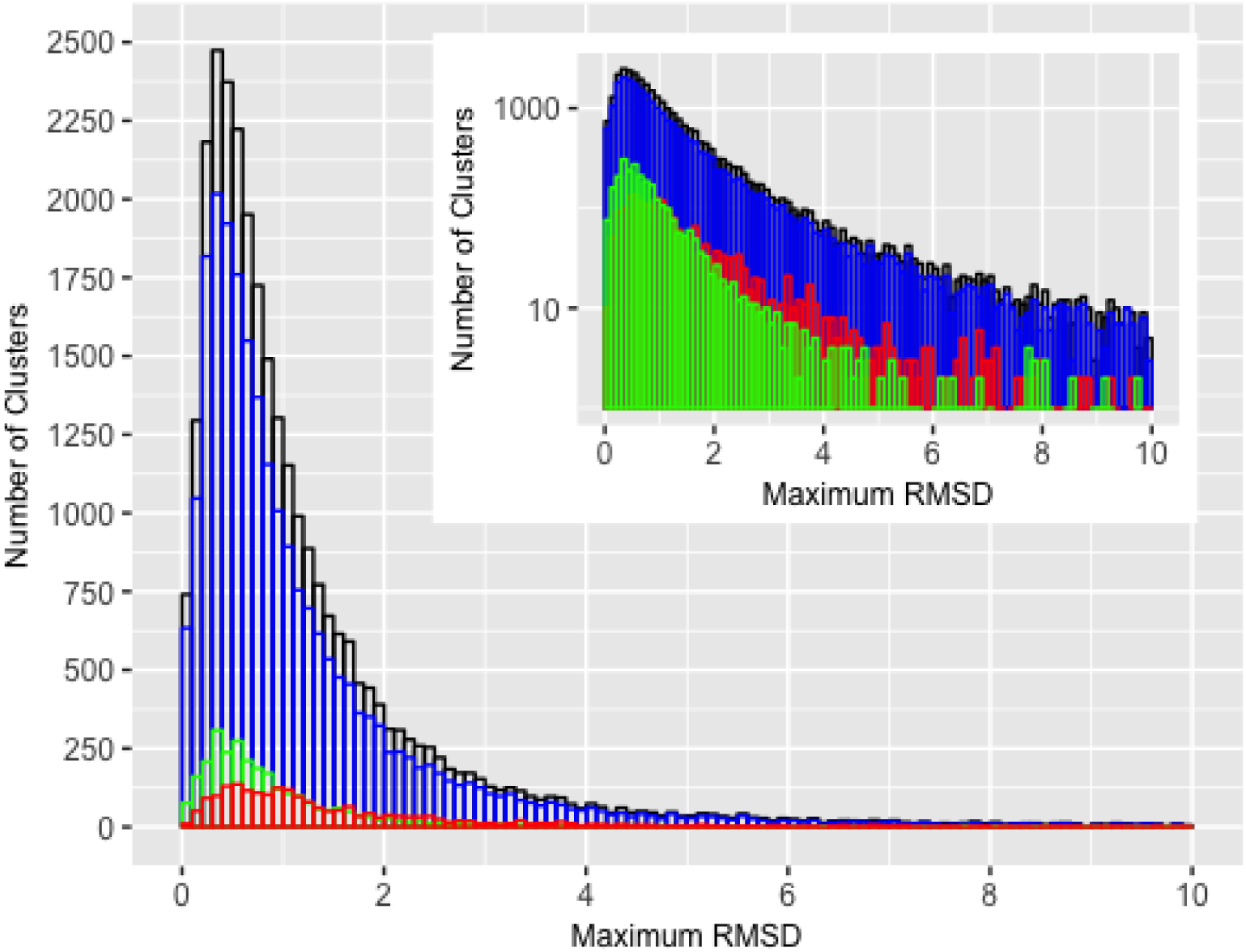
Distribution of maximum RMSD shows the same trend for all clusters with a large peak at 0.4Å and a long tail toward higher RMSDs. *Inset:* The same distribution on a log scale. Black represents the distribution for all clusters, blue represents clusters with two ligand or complex -bound structures, red represents structures with one bound and one unbound structure, and green represents clusters with two unbound structures. The maximum RMSD data from the PDBFlex and the data on complex formation from the PDB. The ligand binding information was obtained from the BioLiP database [1].

This picture is qualitatively similar to one presented in a previous analysis from our group [13], showing that despite doubling of the number of entries in the PDB and significant differences in the protocol between the two papers, the general observation of the PDB providing information on the conformational ensemble of proteins remains the same. It is important to note that our analysis provides the lower estimate of the structural divergence for every protein, since introduction of new structures that may be determined in the future can make the maximum RMSD only higher.

### Binding events are not the only cause for conformational changes

In order to determine to what extent the structural changes between alternative conformations were caused by ligand binding/complex formation, we classified the clusters into three classes, based on whether one or both of the structures in the maximum RMSD pair were or were not bound to a ligand or part of a complex. If binding or complex formation were the only causes for conformation change, then only the clusters represented by one ligand (or complex) bound structure and one free structure would show significant differences in their RMSD. However, as seen in Figure 2, we found that the distribution of RMSD values was similar for all three classes, with a long tail towards higher RMSD values, indicating that conformation changes are not always caused by binding or complex formation alone. This result is in line with several studies that have shown that ligand binding or complex formation does not “induce” structural changes, but results in the stabilization of one of many possible conformations and a shift in the conformational equilibrium towards that conformation [21, 22].

To assess the contribution of different crystallization conditions/experiments to the structural differences, we classified the clusters into two groups based on whether the two most divergent structural models in the cluster were different chains/coordinate sets in the same PDB entry (representing the same crystallization experiment) or were from different PDB entries (and would therefore be likely to represent different experimental conditions). We found that while overall the two distributions looked similar, the maxima were shifted by 0.2 Å and the ratio between both groups reversed between the small and larger RMSD values (See Figure 3). For instance, the most divergent pair in the ∼66% of the clusters (6000/9079 clusters) with RMSD ≤ 0.5Å came from the same PDB entry. For clusters with RMSD > 0.5Å, this ratio was reversed with ~46% of clusters (9674/20934 clusters) represented by models from the same PDB entries (Figure 3 inset). These results show that different crystallization experiments can explore a broader range of conformational flexibility of proteins and further underscores the observation that the redundancy in the PDB can be used to study protein conformational diversity.

**Figure 3.**
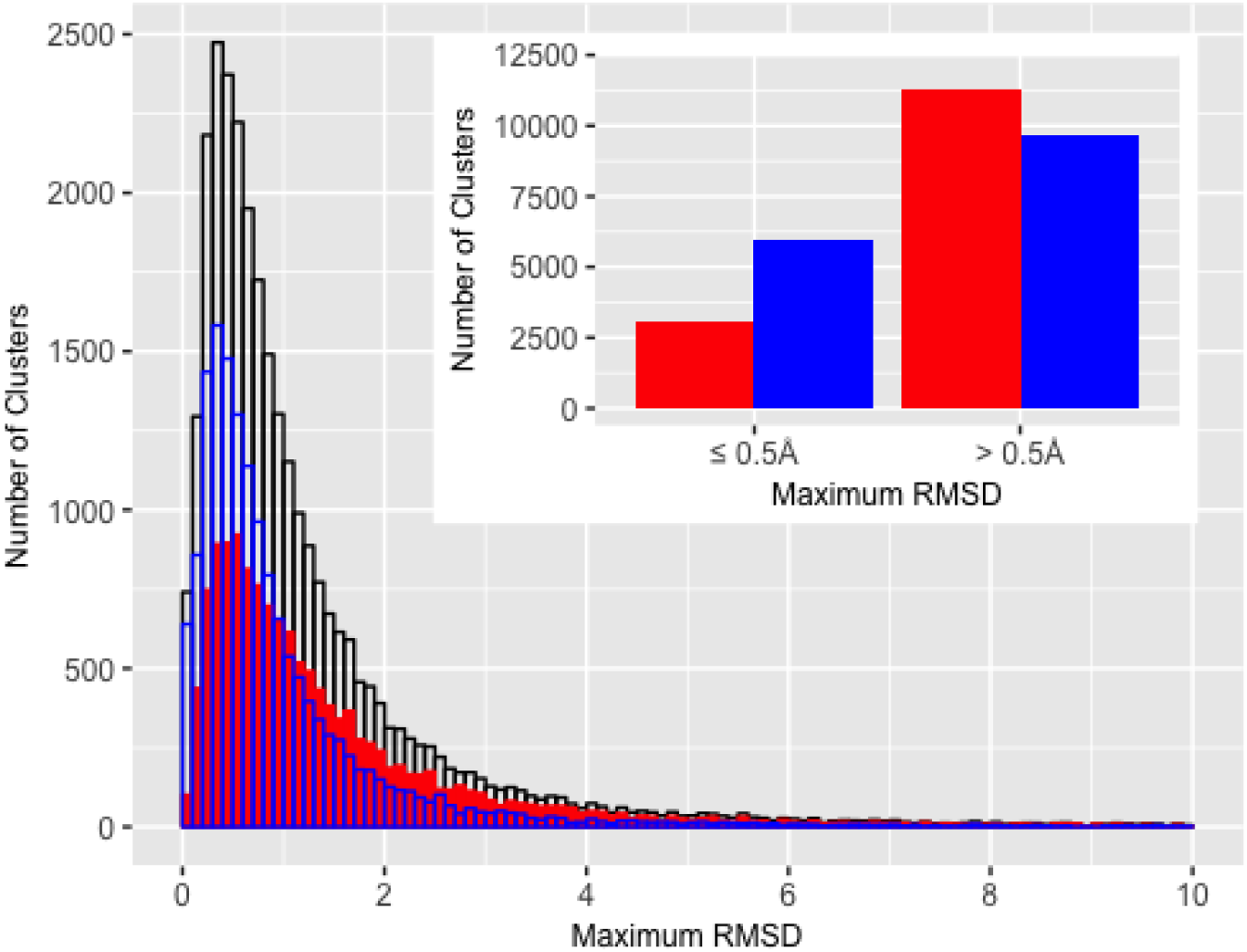
Distribution of maximum RMSD for all clusters (black), clusters represented by structural models/chains belonging to the same PDB entry (blue) or to different entries (red). Inset: ratio between red and blue groups reverses between low and high RMSD.

We hypothesized that differential contacts could be modulating the equilibrium between alternative conformations seen in the PDB. To verify this, we looked at the fraction of the total number of contacts that have changed between two conformations (as seen in the filtered differential contact maps) for three classes of clusters: those represented by one bound and one free (or holo- and apo-)) structure (Figure 4). In order to see some examples of which residues are identified as the DSRs, we applied this approach to two examples (presented below) of proteins that bind substrates/form complexes

**Figure 4.**
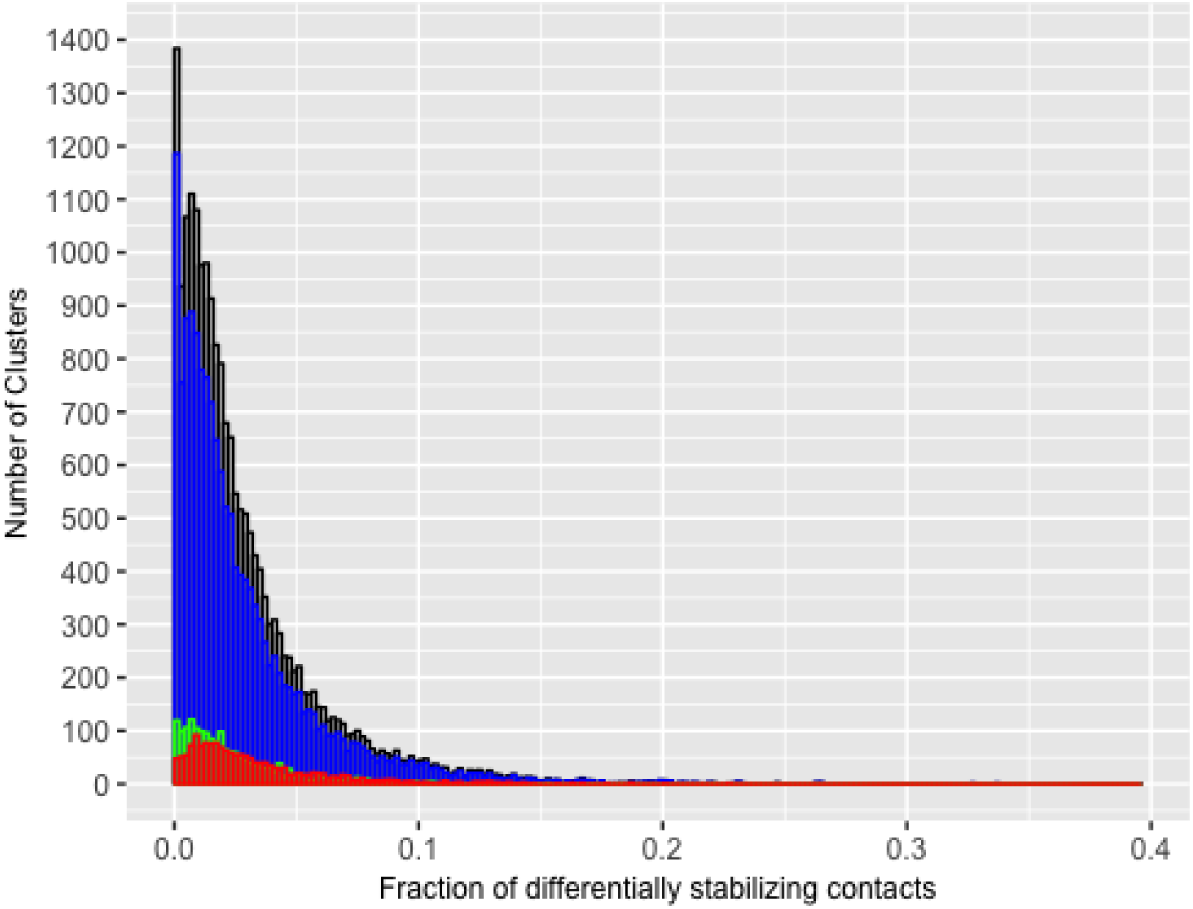
The distribution of the fraction of flexibility modulating contacts for all clusters. Black represents the distribution for all clusters, blue represents clusters with two ligand-bound structures, red represents structures with one bound and one unbound structure, and green represents clusters with two unbound structures. This distribution has a similar shape for all categories, with a long right-tail.

### Adenylate Kinase

Adenylate kinase (ADK) is an enzyme that catalyze the transfer of one phosphate group from ATP to AMP, to form two ADP molecules. ADK consists of three domains: the NMP_bind_ domain, the LID domain and the CORE domain [23]. Substrate-binding and catalysis by this enzyme involves a conformation change wherein the NMP_bind_ and LID domains move from an “open” to a “closed” conformation. However, this enzyme has been known to sample both conformations even in the absence of the substrate [24, 25].

We created contact maps for the closed and open conformations based on the PDB models with codes 4jzkB and 4akeA [23], respectively. The final differential contact map showed a total of 49 contacts unique to only one conformation, 31 to the closed form 18 to the open form (Figure 5). Six contacts were seen between the NMP_bind_ and LID domains, nine between the NMP_bind_ and CORE domains and eight contacts were between the LID and CORE domains. The remaining contacts were intra-domain contacts. We hypothesized that the inter-domain contacts would be responsible for modulating the conformational change seen in this protein. One of the six contacts between the NMP_bind_ and CORE domains was formed by Glu170 and Leu58 and seen only in the closed conformation. It has been previously shown that a hydrogen bond exists between these residues in the closed conformation and that the disruption of this bond (by mutation of Glu170 to Alanine) resulted in a shift of the conformational equilibrium towards the open form in the absence of the substrate [24]. In order to assess whether this mutation could be impacting the differential stability of the two conformations, we used MODELLER [26] to build two structures for the WT sequence and two structures for the mutated sequence, using 4akeA and 4jzkB as templates. We then calculated the DOPE scores for these modeled structures and found that the open conformation had a higher DOPE score for both the WT and mutated structures (Table 1). However, the difference in the scores for the open and closed conformation was less in the mutant structure (Table 1), which suggests a shift in the equilibrium to the open form [24].

**Figure 5.**
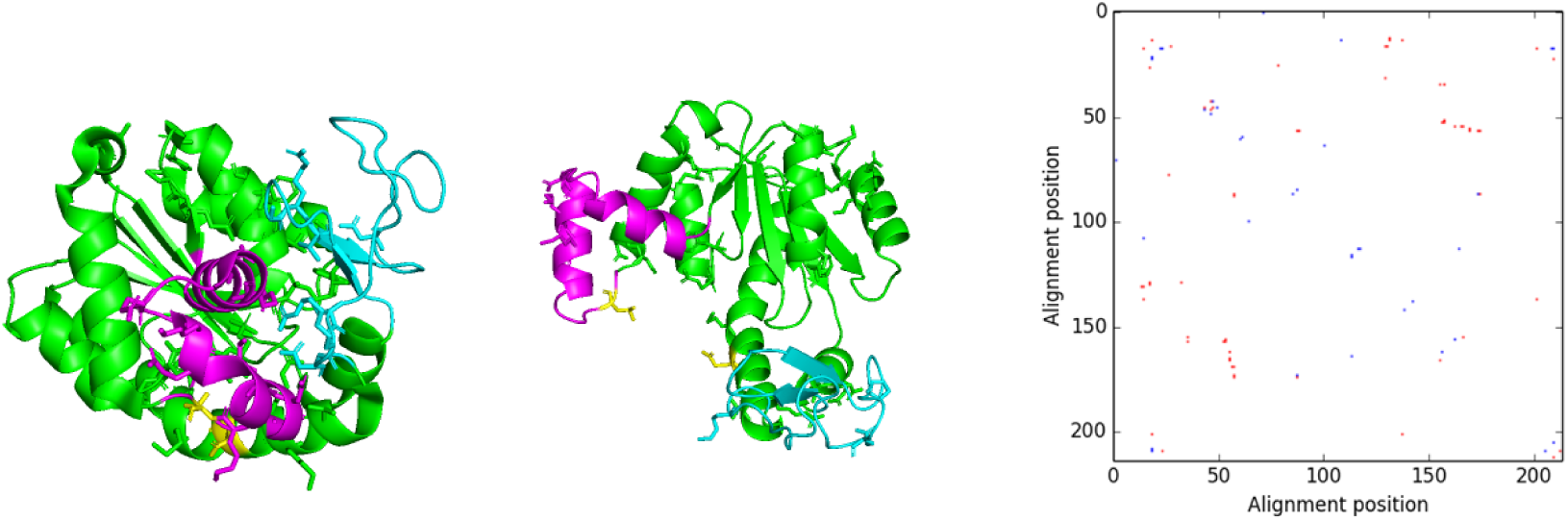
DSRs seen in adenylate kinase. The cartoon representations of the (a) closed and (b) open forms of the ADK. The NMP_bind_ domain is colored magenta, the LID domain is cyan and the CORE domain is green. Residues forming contacts are shown as sticks. Glu170 and Leu58 are shown in yellow sticks in both structures. (c) The final differential contact map for this enzyme. Red contacts are seen in the closed conformation and blue contacts are seen in the open conformation.

**Table 1.**
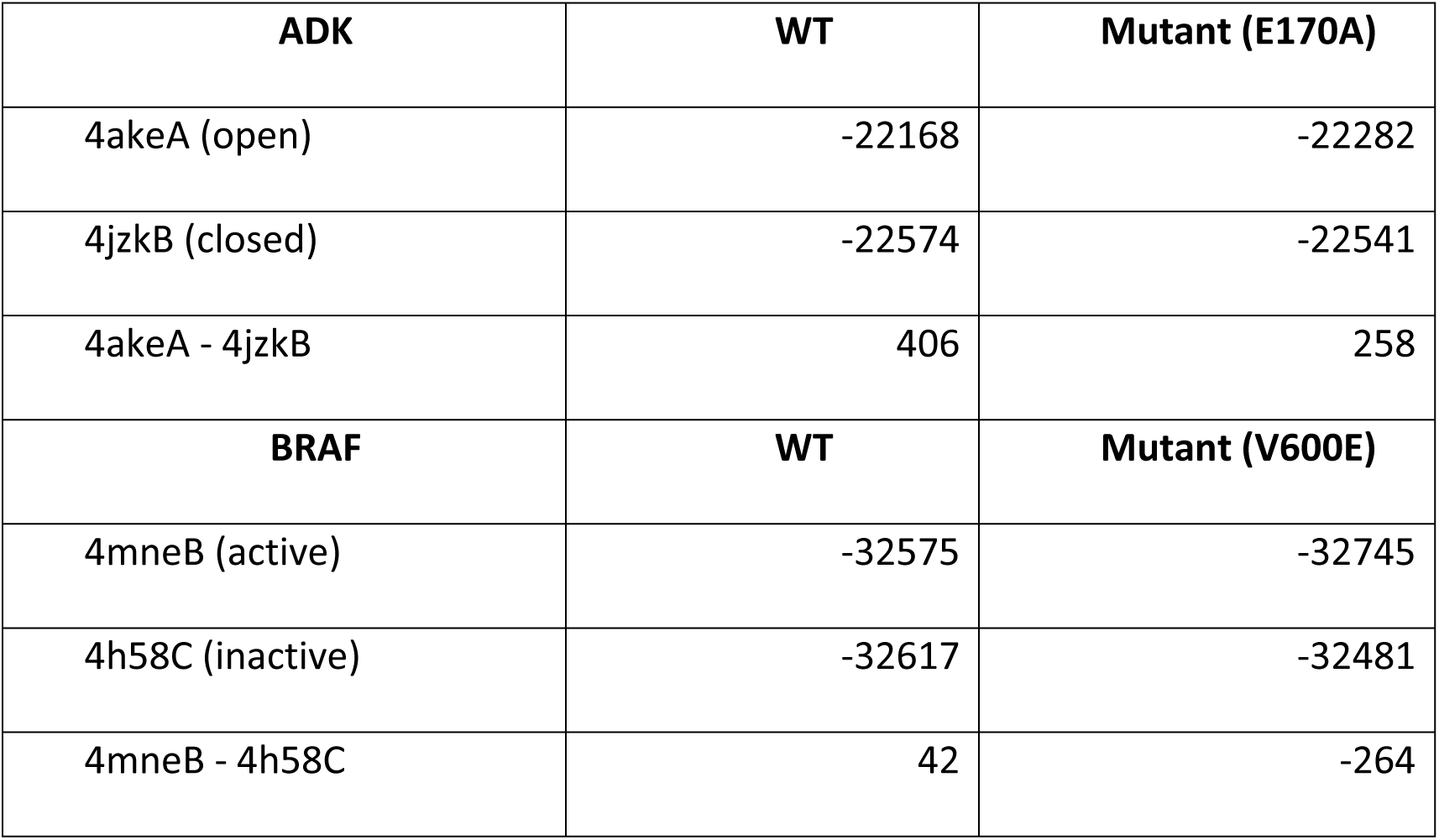
DOPE scores for the modeled WT and mutant conformations of ADK and BRAF. The difference in the DOPE scores for the two ADK conformations suggests that the mutation results in a shift towards the open conformation, while for BRAF, the scores suggest a stabilization and shift of the equilibrium towards the active conformation.

| **ADK** | **WT** | **Mutant (E170A)** |
| --- | --- | --- |
| 4akeA (open) | -22168 | -22282 |
| 4jzkB (closed) | -22574 | -22541 |
| 4akeA - 4jzkB | 406 | 258 |
| **BRAF** | **WT** | **Mutant (V600E)** |
| 4mneB (active) | -32575 | -32745 |
| 4h58C (inactive) | -32617 | -32481 |
| 4mneB - 4h58C | 42 | -264 |

### BRAF

BRAF is a protein kinase and one of the components of the Ras-Raf-MEK-ERK signaling pathway that plays a central role in cell proliferation and differentiation. Activation of BRAF allows it to phosphorylate MEK via its kinase domain [27]. The BRAF kinase domain consists of a small N-terminal lobe and a large C-terminal lobe. The N-terminal lobe includes the glycine rich P-loop (residues 463-471) and has a major α-helix termed the αC-helix. The C-terminal lobe includes the activation segment/AS beginning with the DFG motif (residues 594-596) and ending with the APE motif (residues 621-623) [28].

We created contact maps for BRAF based on the models with PDB codes 4mneB (in which it is complexed to MEK1) [29] and 4h58C (in which BRAF is bound to an inhibitor) [30]. The final differential contact map showed 82 contacts, 51 of which are between residues from the activation segment and the rest of the protein. This included the contact between R575 and E611, that has been described to stabilize the active conformation of the protein [29].

Given its role in cell growth and division, BRAF is frequently mutated in many different cancers. One of the most common mutations seen in this protein is the V600E mutation, that accounts for 80% of all BRAF mutations [31]. This residue forms a contact with K507 that is responsible for providing additional stability to the active conformation in the mutant form of the protein [29]. This contact was seen in our final differential contact map, demonstrating a possible application of this method to exploring the role of various disease-causing mutations.

## Discussion

In this manuscript we have explored a contact map based description of protein structures with a goal of identifying residues that preferentially stabilize alternative conformations, potentially controlling the balance between different functional isoforms. Exploring the redundancy in the Protein Data Bank we realized that, for most proteins, multiple coordinate sets deposited in PDB provide some level of sampling of these proteins’ conformational ensemble. We have shown that while for most proteins the differences between various coordinate sets available in the PDB is small, for a significant number of them the differences are not trivial and, most likely, sample different functional states. Somewhat surprisingly, these differences are not necessarily related to ligand binding and complex formation and even models of unliganded proteins solved in different crystal forms provide a significant amount of information about the shape of the conformational ensemble.

We developed an automated protocol to identify residues involved in stabilizing alternative conformations of proteins and integrated this information with the PDBFlex database, previously developed in our group. As PDBFlex is integrated directly with the PDB database and website, this provides information about such differentially stabilizing residues (DSRs) for hundreds of proteins to a broad group of users. We explored the DSRs identified by our protocol on few examples of clinically important proteins and confirmed that they indeed overlap with residues overlap with residues previously identified to stabilize alternative conformation and/or mutated in disease states.

Differential contact map visualization and identification of DSRs is now implemented on the PDBFlex server, with thousands of examples open for user analysis. We believe that this resource can be used as a hypothesis-generation tool to identify the residues that could be modulating the conformational equilibrium of any given protein.

## Materials and Methods

### Calculation of Contact Maps and Identification of DSRs

We have developed a method based on contact maps to identify residues that could be modulating the conformational flexibility of a protein by forming contacts that differentially stabilize two different conformations of the protein. This method is described below and illustrated by figure 7.

**Figure 6.**
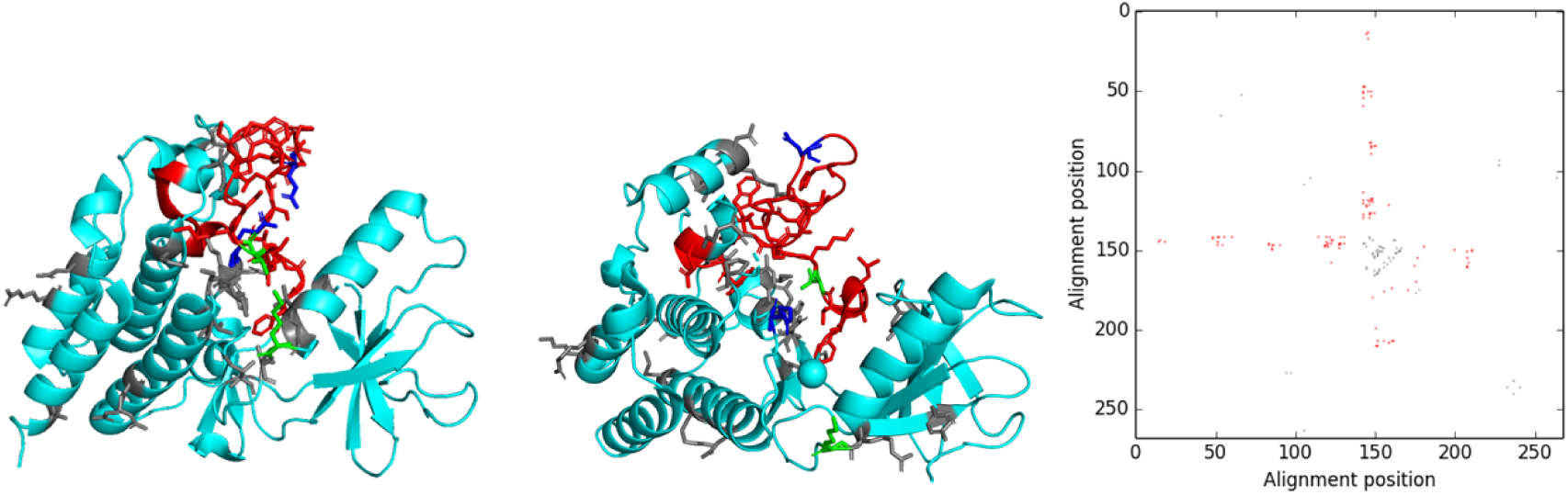
DSRs seen in BRAF. Cartoon representations of BRAF and the differential contact map showing differentially stabilizing contacts. The two structures with maximum RMSD are represented by (a) 4mneB and (b) 4h58C. The residues involved in flexibility modulating contacts are shown represented by sticks. Residues in red form the activation segment. Residues R575 and E611 are in blue, while V600 and K507 are in green. (c) The final differential contact map representing all DSRs. Red contacts represent those formed between activation segment residues and other parts of the protein.

**Figure 7.**
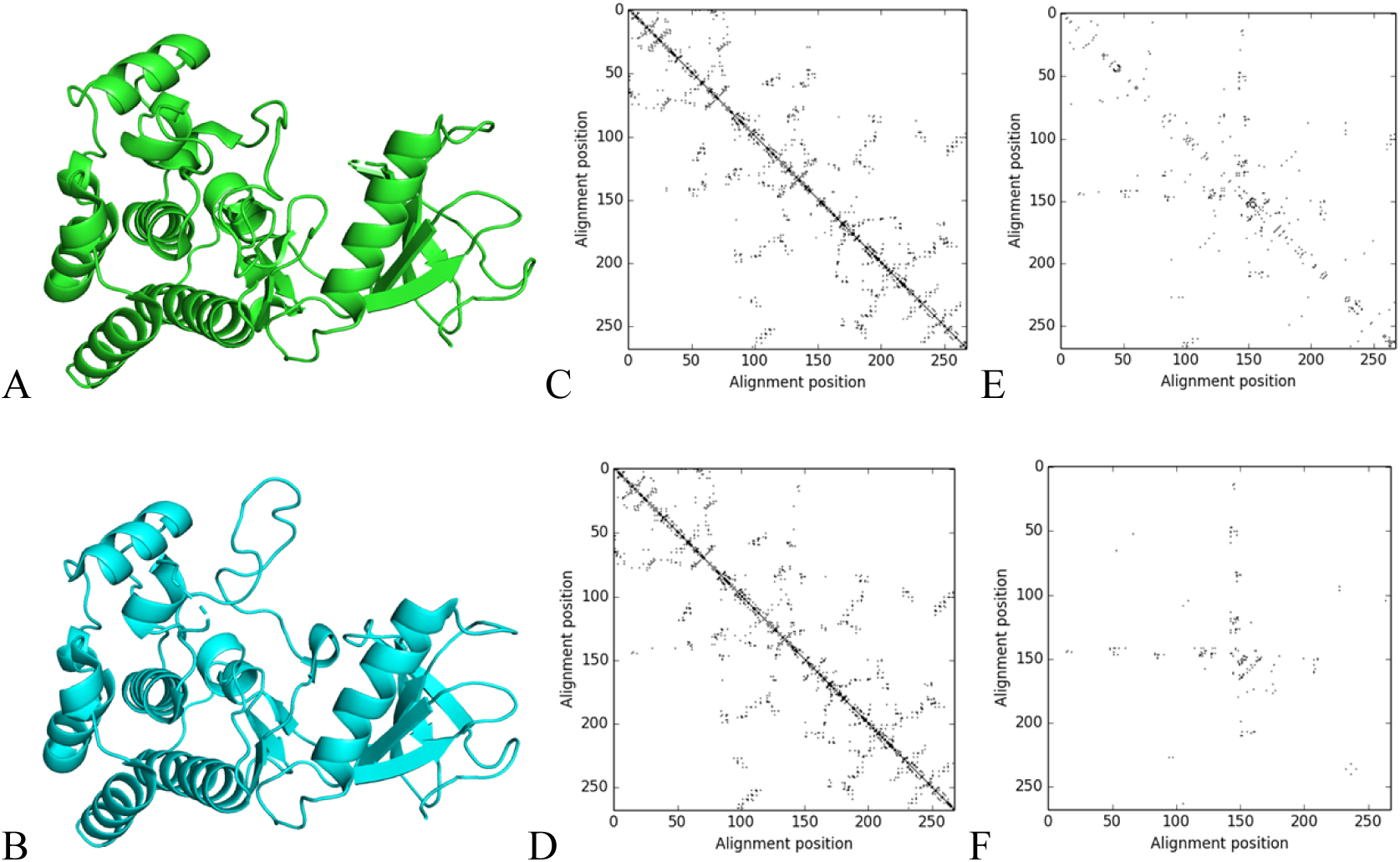
Intermediate stages of the pipeline used for identifying differentially stabilizing residues in BRAF. A) The active and (B) inactive conformations of BRAF in cartoon form (PDB codse: 4mneB and 4h58C respectively). Contact maps for the (C) active and (D) inactive conformations. (E) The differential contact map obtained by subtracting C and D. (F) The final differential contact map obtained after filtering e.

#### Selection of structural models and alignment of sequences

We used structural models deposited in the Protein Data Bank clustered at 95% sequence identity, as done in PDBFlex [14]. Each cluster represents various conformations of a single protein (within the 95% sequence identity threshold). For each cluster, we selected the pair of structural models with maximum RMSD as representative of the two most divergent conformations of the protein (Figure 7a and 7b). However, this method could be performed for any two structural models of the protein. The sequence alignment for the two structural models, based on the PDB SEQRES sequences, was modified to remove any residues that did not have coordinates in the model. This alignment was further modified to remove any residues that were aligned to a gap in either of the sequences.

#### Calculation of contact maps

For each structural model, PDB files were downloaded and the coordinates of the residues in the alignment, corrected as described above were extracted. Only residues with an empty insertion code and hetero-flag were used. (If more than 5% of the extracted residues differed between the two structural models, or the total number of residues extracted was not the same, then the cluster was skipped). Based on the coordinates for these residues, contact maps were created for the two structural models (Figure 7c and 7d). A pair of residues was considered to be in contact if the distance between any two side-chain heavy atoms from the residues was less than 5Å. Residues that did not have coordinates for any heavy atoms were considered to not form contacts and for glycines, the Cα atom was used.

#### Calculation and filtering of differential contact maps

A differential contact map was then created by subtracting one contact map from the other. This map therefore, represents the set of contacts seen in only one conformation, but not the other (Figure 7e). We further filtered this differential contact map to remove any residue pairs that showed less than 5Å difference in inter-residue distance in the two conformations (Figure 7f). This was done in order to consider only the residues that could be modulating large scale conformation changes and to exclude contact changes seen as a result of small local fluctuations in the structure. The residues forming the contacts seen in this final differential contact map were considered to be differentially stabilizing residues (DSRs) that could be modulating the change between the two conformations considered.

Calculation of DOPE scores: MODELLER v 9.21 [26, 32] was used to create structure models for the WT and mutated sequences of BRAF and ADK, based on template structures representing the two alternative conformations of each protein. Alignments between the sequence and the template structure were created using MODELLER’s align2d function. Models were created using the automodel class, with the DOPE score as the assessment method. One model was created per template, for each sequence.

For adenylate kinase, the SEQRES sequence for chain A from the PDB file for 4ake was used to create a sequence file in the MODELLER .ali format. This sequence was manually edited to create a sequence corresponding to the E170A mutation. The WT and mutated sequence were modeled using 4akeA and 4jzkB as templates.

For BRAF, the SEQRES sequence for chain C from the PDB file for 4h58C was used. 4mneB and 4h58C were used as template structures. Since the PDB ATOM records for both 4mneB and 4h58C start from residue D449 (D448 in 4h58C), the sequence file was manually edited to start the sequence from the same residue and reduce the number of gaps in the resulting alignment. The same procedure was followed for the V600E mutated sequence.

## Code and data availability

All the supplementary tables, the raw data and the algorithms used in this manuscript, as well as the results presented, can be downloaded from the PDBFlex server.

## References

1. Yang J, Roy A, Zhang Y. BioLiP: a semi-manually curated database for biologically relevant ligand-protein interactions. Nucleic Acids Res. 2013;41(Database issue):D1096–103. Epub 2012/10/23. doi:10.1093/nar/gks966. PubMed PMID: 23087378; PubMed Central PMCID: PMCPMC3531193.

2. Richardson JS. Schematic drawings of protein structures. Methods Enzymol. 1985;115:359–80. Epub 1985/01/01. PubMed PMID: 3853075.

3. Schrodinger, LLC. The PyMOL Molecular Graphics System, Version 1.8. 2015.

4. Westhead DR, Slidel TW, Flores TP, Thornton JM. Protein structural topology: Automated analysis and diagrammatic representation. Protein Sci. 1999;8(4):897–904. Epub 1999/04/22. doi:10.1110/ps.8.4.897. PubMed PMID: 10211836; PubMed Central PMCID: PMCPMC2144300.

5. Phillips DC. The development of crystallographic enzymology. Biochem Soc Symp. 1970;30:11–28. Epub 1970/01/01. PubMed PMID: 4923824.

6. Godzik A, Skolnick J, Kolinski A. Regularities in interaction patterns of globular proteins. Protein Eng. 1993;6(8):801–10. Epub 1993/11/01. PubMed PMID: 8309927.

7. Cooper A. Thermodynamic fluctuations in protein molecules. Proc Natl Acad Sci U S A. 1976;73(8):2740–1. Epub 1976/08/01. PubMed PMID: 1066687; PubMed Central PMCID: PMCPMC430724.

8. Berman HM, Westbrook J, Feng Z, Gilliland G, Bhat TN, Weissig H, et al. The Protein Data Bank. Nucleic Acids Res. 2000;28(1):235–42. Epub 1999/12/11. PubMed PMID: 10592235; PubMed Central PMCID: PMCPMC102472.

9. Maiorov VN, Crippen GM. Significance of root-mean-square deviation in comparing three-dimensional structures of globular proteins. J Mol Biol. 1994;235(2):625–34. Epub 1994/01/14. doi:10.1006/jmbi.1994.1017. PubMed PMID: 8289285.

10. Zhang Y, Skolnick J. TM-align: a protein structure alignment algorithm based on the TM-score. Nucleic Acids Res. 2005;33(7):2302–9. Epub 2005/04/26. doi:10.1093/nar/gki524. PubMed PMID: 15849316; PubMed Central PMCID: PMCPMC1084323.

11. Nishikawa K, Ooi T. Comparison of homologous tertiary structures of proteins. J Theor Biol. 1974;43(2):351–74. Epub 1974/02/01. PubMed PMID: 4818352.

12. Wang G, Dunbrack RL, Jr. PISCES: a protein sequence culling server. Bioinformatics. 2003;19(12):1589–91. Epub 2003/08/13. PubMed PMID: 12912846.

13. Burra PV, Zhang Y, Godzik A, Stec B. Global distribution of conformational states derived from redundant models in the PDB points to non-uniqueness of the protein structure. Proc Natl Acad Sci U S A. 2009;106(26):10505–10. Epub 2009/06/26. doi:10.1073/pnas.0812152106. PubMed PMID: 19553204; PubMed Central PMCID: PMCPMC2705611.

14. Hrabe T, Li Z, Sedova M, Rotkiewicz P, Jaroszewski L, Godzik A. PDBFlex: exploring flexibility in protein structures. Nucleic Acids Res. 2016;44(D1):D423–8. Epub 2015/11/29. doi:10.1093/nar/gkv1316. PubMed PMID: 26615193; PubMed Central PMCID: PMCPMC4702920.

15. Monzon AM, Rohr CO, Fornasari MS, Parisi G. CoDNaS 2.0: a comprehensive database of protein conformational diversity in the native state. Database (Oxford). 2016;2016. Epub 2016/03/30. doi:10.1093/database/baw038. PubMed PMID: 27022160; PubMed Central PMCID: PMCPMC4809262.

16. Andonov R, Djidjev H, Klau GW, Boudic-Jamin ML, Wohlers I. Automatic Classification of Protein Structure Using the Maximum Contact Map Overlap Metric. 2015;8(4):850. PubMed PMID: doi:10.3390/a8040850.

17. Vehlow C, Stehr H, Winkelmann M, Duarte JM, Petzold L, Dinse J, et al. CMView: interactive contact map visualization and analysis. Bioinformatics. 2011;27(11):1573–4. Epub 2011/04/08. doi:10.1093/bioinformatics/btr163. PubMed PMID: 21471016.

18. Andonov R, Malod-Dognin N, Yanev N. Maximum contact map overlap revisited. J Comput Biol. 2011;18(1):27–41. Epub 2011/01/08. doi:10.1089/cmb.2009.0196. PubMed PMID: 21210730.

19. Liu L, Gronenborn AM, Bahar I. Longer simulations sample larger subspaces of conformations while maintaining robust mechanisms of motion. Proteins. 2012;80(2):616–25. Epub 2011/11/23. doi:10.1002/prot.23225. PubMed PMID: 22105881; PubMed Central PMCID: PMCPMC3290687.

20. Go N, Noguti T, Nishikawa T. Dynamics of a small globular protein in terms of low-frequency vibrational modes. Proc Natl Acad Sci U S A. 1983;80(12):3696–700. Epub 1983/06/01. PubMed PMID: 6574507; PubMed Central PMCID: PMCPMC394117.

21. Boehr DD, Nussinov R, Wright PE. The role of dynamic conformational ensembles in biomolecular recognition. Nat Chem Biol. 2009;5(11):789–96. Epub 2009/10/21. doi:10.1038/nchembio.232. PubMed PMID: 19841628; PubMed Central PMCID: PMCPMC2916928.

22. Wei G, Xi W, Nussinov R, Ma B. Protein Ensembles: How Does Nature Harness Thermodynamic Fluctuations for Life? The Diverse Functional Roles of Conformational Ensembles in the Cell. Chem Rev. 2016;116(11):6516–51. Epub 2016/01/26. doi:10.1021/acs.chemrev.5b00562. PubMed PMID: 26807783.

23. Muller CW, Schlauderer GJ, Reinstein J, Schulz GE. Adenylate kinase motions during catalysis: an energetic counterweight balancing substrate binding. Structure. 1996;4(2):147–56. Epub 1996/02/15. PubMed PMID: 8805521.

24. Aden J, Verma A, Schug A, Wolf-Watz M. Modulation of a pre-existing conformational equilibrium tunes adenylate kinase activity. J Am Chem Soc. 2012;134(40):16562–70. Epub 2012/09/12. doi:10.1021/ja3032482. PubMed PMID: 22963267.

25. Arora K, Brooks CL, 3rd. Large-scale allosteric conformational transitions of adenylate kinase appear to involve a population-shift mechanism. Proc Natl Acad Sci U S A. 2007;104(47):18496–501. Epub 2007/11/15. doi:10.1073/pnas.0706443104. PubMed PMID: 18000050; PubMed Central PMCID: PMCPMC2141805.

26. Sali A, Blundell TL. Comparative protein modelling by satisfaction of spatial restraints. J Mol Biol. 1993;234(3):779–815. Epub 1993/12/05. doi:10.1006/jmbi.1993.1626. PubMed PMID: 8254673.

27. Robinson MJ, Cobb MH. Mitogen-activated protein kinase pathways. Curr Opin Cell Biol. 1997;9(2):180–6. Epub 1997/04/01. PubMed PMID: 9069255.

28. Roskoski R, Jr. RAF protein-serine/threonine kinases: structure and regulation. Biochem Biophys Res Commun. 2010;399(3):313–7. Epub 2010/08/03. doi:10.1016/j.bbrc.2010.07.092. PubMed PMID: 20674547.

29. Haling JR, Sudhamsu J, Yen I, Sideris S, Sandoval W, Phung W, et al. Structure of the BRAF-MEK complex reveals a kinase activity independent role for BRAF in MAPK signaling. Cancer Cell. 2014;26(3):402–13. Epub 2014/08/27. doi:10.1016/j.ccr.2014.07.007. PubMed PMID: 25155755.

30. Vasbinder MM, Aquila B, Augustin M, Chen H, Cheung T, Cook D, et al. Discovery and optimization of a novel series of potent mutant B-Raf(V600E) selective kinase inhibitors. J Med Chem. 2013;56(5):1996–2015. Epub 2013/02/13. doi:10.1021/jm301658d. PubMed PMID: 23398453.

31. Davies H, Bignell GR, Cox C, Stephens P, Edkins S, Clegg S, et al. Mutations of the BRAF gene in human cancer. Nature. 2002;417(6892):949–54. Epub 2002/06/18. doi:10.1038/nature00766. PubMed PMID: 12068308.

32. Webb B, Sali A. Comparative Protein Structure Modeling Using MODELLER. Curr Protoc Protein Sci. 2016;86:2 9 1–2 9 37. Epub 2016/11/02. doi:10.1002/cpps.20. PubMed PMID: 27801516.

